# Decoding protein binding landscape on circular RNAs with base-resolution Transformer models

**DOI:** 10.1101/2022.11.20.517239

**Authors:** Hehe Wu, Yi Fang, Yang Yang, Xiaoyong Pan, Hong-Bin Shen

## Abstract

Circular RNAs (circRNAs) interact with RNA-binding proteins (RBPs) to modulate gene expression. To date, most computational methods for predicting RBP binding sites on circRNAs focus on circRNA fragments instead of circRNAs. These methods detect whether a circRNA fragment contains binding sites, but cannot determine where are the binding sites and how many binding sites are on the circRNA transcript. We report a hybrid deep learning-based tool, CircSite, to predict RBP binding sites at single-nucleotide resolution and detect key contributed nucleotides on circRNA transcripts. CircSite takes advantage of convolutional neural networks (CNNs) and Transformer for learning local and global representations of circRNAs binding to RBPs, respectively. We construct 37 datasets of RBP-binding circRNAs for benchmarking and the experimental results show that CircSite offers accurate predictions of RBP binding nucleotides and detects key subsequences aligning well with known binding motifs.

## 1. Introduction

Circular RNAs (circRNAs) interact with RNA-binding proteins (RBPs) to modulate gene expression [1] and accomplish biological functions. For instance, CircRNA ciRS-7 functions as miRNA-7 sponge by binding to AGO protein [2]. Meanwhile, RBPs play crucial roles in many biological processes. For CRISPR/Cas9 genome editing technique, a guide RNA (gRNA) binds to the Cas9 protein to modulate it, activating the endonuclease activity on the DNA targets [3]. Protein PEG10 binds and packages RNAs as an RNA delivery tool [4]. Furthermore, the interactions between circCDYL and RBPs affect the cancer pathway associated with bladder cancer [5]. Thus, identifying the interactions between circRNAs and RBPs provides insights into the functions of RBPs and circRNAs, further revealing the mechanism behind diseases [6].

With the development of high-throughput sequencing technologies [7], many binding targets including circRNAs of RBPs have been collected [8, 9]. Based on the accumulated binding targets, RBP-specific machine learning approaches have been proposed to predict RBP binding sites on linear and circular RNAs [10–24] and they generally train a model per RBP. For linear RNAs, GraphProt first encodes the sequence and predicted structure to a graph, which is fed into support vector machine for RBP-binding site classification [25]. iONMF uses orthogonal matrix factorization to integrate multiple sources of features to predict RBP binding sites [26]. DeepBind [10] trains convolutional neural networks (CNNs) to infer the binding sequence preference of RBPs. iDeepS [27] further applies hybrid CNNs and long short-term memory neural network (LSTMs) to learn binding sequence and structure preference of RBPs simultaneously, where LSTMs capture long dependence between sequences and structures. Similarly, DeepCLIP also trains a hybrid CNN and LSTM model to predict the impact of mutations on RNA-protein interactions [12]. For circRNAs, CRIP designs a stacked codon-based encoding scheme with a hybrid network of CNNs and LSTMs to predict RBP binding sites on circRNAs [14]. PASSION designs an ensemble neural network to identify RBP binding sites on circRNAs [28]. iCircRBP-DHN constructs a deep network for predicting RBP binding sites on circRNAs using embedding and K-tuple nucleotide frequencies [16]. iDeepC applies Siamese network to handle the poorly characterized RBPs with a few of binding circRNA targets [29]. In addition, the CNN-based models can extract the binding motifs from the learned kernels of CNNs, but these motifs cannot provide nucleotide-level interpretation. Since the training of deep models is time-consuming, some easy-of-use online webservers, i.e. RBPsuite [17] that implements CRIP and iDeepS as online service and DeepCLIP [12], are developed.

In general, the above methods only provide relatively low-resolution binding regions at a predefined number of nucleotides. They mainly focus on predicting whether an RNA fragment contains RBP binding sites or not, where these fragments are subsequences centering at the binding sites or nucleotides. For example, GraphProt constructs the training fragments by extending the binding sites in downstream and upstream regions by 150 nucleotides [30]. iONMF collects the training fragments with a fixed length of 101-bp by extending the binding nucleotides in both directions by 50 nucleotides [26]. However, the models trained on these data focus on predicting the enriched binding regions at about 100-bp resolution and could not precisely locate the binding nucleotides on the circRNAs. To apply the trained models to circRNAs, we can first use a fixed-size sliding window to scan the circRNAs with a step one into fragments, then use the trained models to predict binding scores for individual fragments. However, these strategies suffer from a high false positive rate problem [29] and treat each nucleotide independently when making decision, which ignores the binding status of neighboring nucleotides. Thus, it is imperative to develop computational tools to model RBP binding landscape at a nucleotide-level resolution precisely and accurately.

In this study, we design a novel interpretable deep model, called CircSite, to predict the RBP binding landscapes on circRNA transcripts at a nucleotide-level resolution (a nucleotide), further inferring RBP binding circRNA regions (several consecutive nucleotides) and circRNAs (a circRNA transcript). CircSite consists of five main modules: a convolutional neural network (CNN), a Bidirectional gated recurrent unit (BiGRU), a Transformer, a median filtering module, and an interpretable module. The CNN learns local high-level abstract representations of circRNA sequences, followed by the BiGRU to learn long-dependency representations of RNA sequences. The Transformer learns the global attention-based representations of circRNA sequences. Both the local representations learned by CNN-BiGRU and the global representations learned by Transformer represent the patterns of circRNAs binding to RBPs. In addition, CircSite uses a median filter to leverage the binding status of neighboring nucleotides to remove false binding nucleotides, followed by a new score binarization strategy to obtain the final binding nucleotides. In the interpretable module, CircSite applies integrated gradient [31] to identify the key sequence contents that contribute to RBP binding landscapes, providing intuitive nucleotide-level interpretation into the decision-making process of deep models. Finally, the experimental results show that CircSite is superior to state-of-the-art methods for predicting binding nucleotides on circRNAs and captures the key subsequences aligning well with known binding motifs.

## 2. Methods

### 2.1. Benchmark datasets

In this study, we construct 37 benchmark datasets of RBP-binding circRNAs for 37 RBPs, each consists of training and test sets at the nucleotide level. In addition, we construct 37 independent test sets consisting of circRNA transcripts for 37 RBPs, these test sets are used to evaluate the performance of CircSite on predicting whether the circRNA can interact with a given RBP.

#### 2.1.1 Nucleotide-level training and test set

We first construct benchmark datasets of RBP binding nucleotides on circRNAs. Over 120,000 circRNAs sequences for 37 RBPs are extracted from the CircInteractome database[32] (https://circinteractome.nia.nih.gov/). For each RBP, we first split the binding sequences into the training and test set with a ratio of 8:2. Considering that high sequence similarity in the training and test sets may lead to overestimated performance, we use CD-HIT [33] with a similarity threshold of 0.8 to remove redundant sequences in the test sets. In addition, to avoid potential bias caused by sequences that were too short or too long, we only keep those sequences with a length between 200 and 6000, which takes over 90% of all the sequences (Supplementary Table S1). The number of circRNAs in the training and test sets for 37 RBPs are given in Supplementary Table S2, and they are used to generate nucleotide-level training and test sets.

To apply deep models to circRNAs, we use a new strategy to generate positive and negative samples from the RNAs as the nucleotide-level training and test set. We scan the sequences by a sliding window with a step of 1 nt to obtain fragment sequences. If the centering nucleotide of the sliding window is within the binding site, then we consider the fragment to be a positive sample, otherwise a negative sample. Thus, each RNA will generate multiple fragments with nucleotide-wise labels. This strategy has two-fold benefits: 1) By using a sliding window approach, we obtain a sufficient number of positive samples and a large number of negative samples, which brings a huge benefit to RBPs with a relatively small amount of known binding circRNAs. Thus, the deep model can learn patterns from a large number of negative samples, resulting in low false positive rates; 2) Different from existing methods that predict whether a fragment contains binding sites or not, this strategy allows the deep model to predict precisely binding nucleotides of RBPs on circRNAs. In the nucleotide-level training and test set, the samples are binding or non-binding nucleotides of RBPs.

#### 2.1.2 Independent circRNA transcript test set

To better explore the advantages of CircSite, we predict whether the circRNA can interact with a given RBP or not. 37 datasets are collected for 37 RBPs, and each of them consists of 100 positive or negative circRNAs randomly selected from circinteractome databases. The positive samples are circRNAs that have binding sites for a given RBP and do not appear in the training set, while the negative samples are circRNAs that do not have any binding sites with the given RBP in circinteractome. In order to make an objective evaluation, we use CD-HIT [33] with a similarity cutoff of 0.8 to ensure the sequences in the independent test set are not similar to the training set. In this independent test set, each sample is a circRNA.

### 2.2. Overview of CircSite framework for predicting RBP binding nucleotides on circRNAs

In this study, we design a hybrid deep network CircSite (Fig. 1) consisting of 1-D CNN, BiGRU and Transformer to predict RBP binding nucleotides on circRNAs, further inferring RBP binding regions and circRNAs. First, we use a sliding window to scan the circRNAs into fragments with a step size of one, and these fragments are represented as a one-hot encoding matrix. CircSite first divides the fragment with a length of the predefined sliding window size into patches by 1D convolution, obtaining representation maps of each patch. Then, CircSite treats each patch as a word, which is fed into the Encoder of the Transformer and BiGRU, respectively. Here, we only use the encoder of Transformer without the decoder. CNN-BiGRU learns local representations by capturing the dependencies between patches and the Transformer learns global representations using the attention mechanism, where the distance between any two positions in the sequence is reduced to a constant. During the training process, BiGRU and Transformer learn RNA representations separately and each has its own advantages. Thus, we concatenate the embeddings learned by the BiGRU and Transformer as the input of an MLP for the final classification. For each nucleotide on the circRNA, CircSite assigns the binding score of the fragment cantering at this nucleotide to it. Then, the scores of individual nucleotides are pooled and post-processed by using a median filter and score binarization to obtain the binding nucleotides on the circRNAs, further inferring RBP binding labels for regions and circRNAs. CircSite is designed to predict RBP binding nucleotides with sequence contexts and the binding information of neighboring nucleotides.

**Fig. 1.**
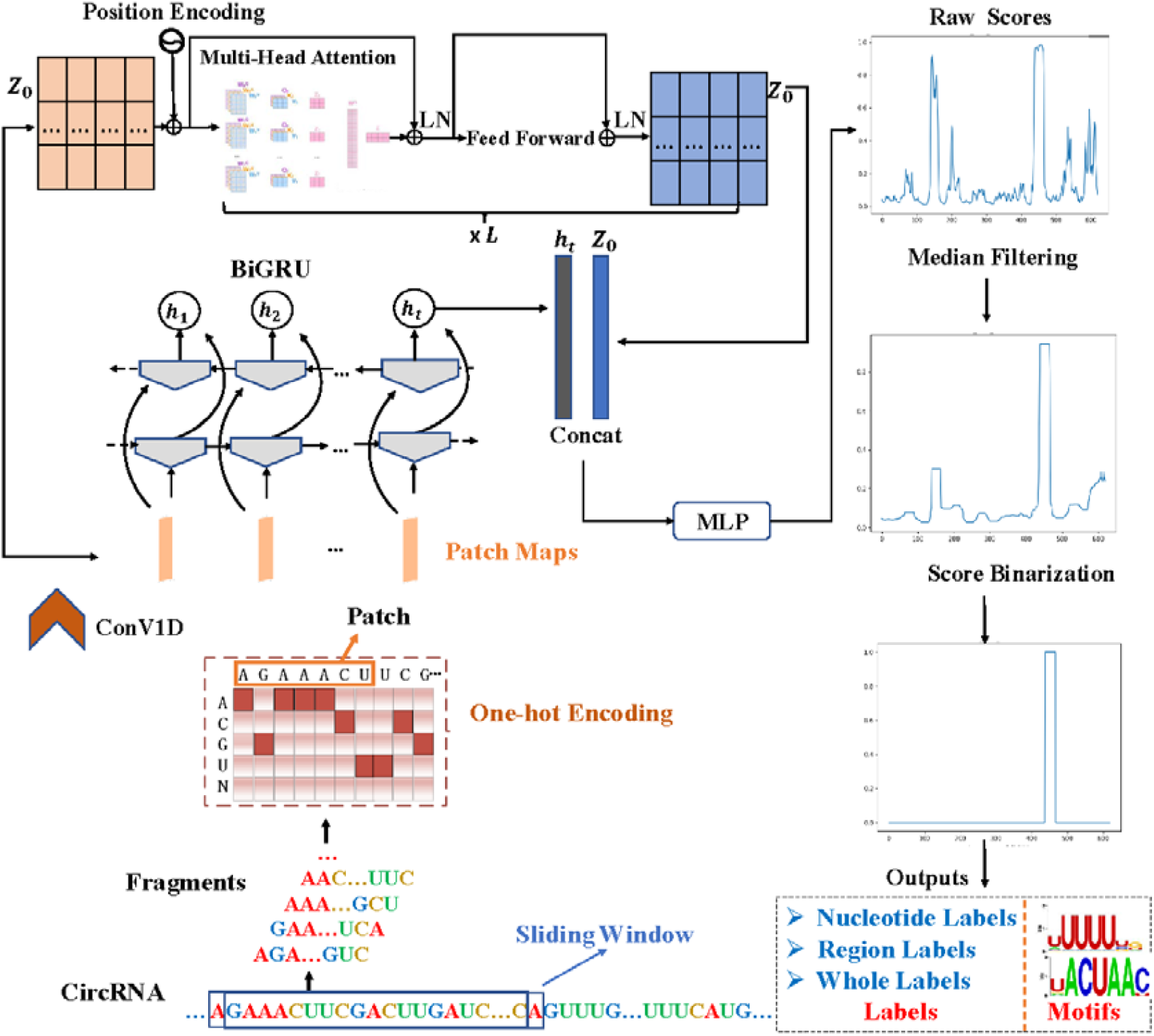
The pipeline of the CircSite. The circRNA is first split into fragments, which are fed into 1-D CNN with a patch input to obtain patch maps, followed by a BiGRU and a Transformer, respectively. Then, the concatenated representations are fed into an MLP classifier, which is used to score each fragment centering at this nucleotide. The raw scores of individual nucleotides on a circRNA are pooled and processed by a median filter and score binarization strategy to obtain the binding nucleotides on the circRNA, further inferring RBP binding labels for circRNA regions and circRNAs. *L* is the number of repetitions of the corresponding block. LN is layer normalization, *h*_*t*_, *Z*_0_ are the output of the last time step of the BiGRU, respectively, and the token is set at dimension 0 of the input matrix.

### 2.3. Convolutional layer

Convolutional neural networks [34] are neural networks with local connectivity and shared weights, which could extract discriminate local information buried in sequences. We first encode a sequence of a length n into a one-hot encoded matrix X of a size *n* * 4, which is fed into a 1-D convolutional layer. We treat a sequence of the convolution kernel size as a patch and map it to a vector as a representation of this patch. Let the size of the convolution kernel be h. Each element 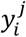in the column vector *y*^*j*^ of the length *n* − *h* + 1 is defined as follow:

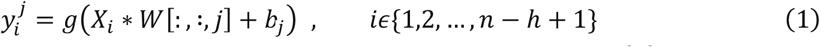

where *g*(·) is the ReLU activation function, *W* is the convolution kernel *W* ∈ *R*^*h*\**k*\*4^ and *b* is the bias.

After this convolutional operation of *k* convolution kernels, the obtained matrix shape is (*B, n* − *h* + 1, *k*), where *B* is the batch size of the input sequence.

### 2.4. BiGRU layer

GRU [35] is a variant of LSTM network but is simpler and more effective. GRU alleviates the problem of gradient vanishing and long-term dependencies by introducing two gates: an update gate and a reset gate. The update gate determines how much information of the old state is copied into the new state and can capture long-term dependency in sequences. The reset gate determines how much information of the old state should be remembered and can capture the short-term dependency in sequences. GRU is more streamlined compared to LSTM. Thus, we choose bidirectional GRU (BiGRU) to explore sequence information.

### 2.5. Transformer layer

The Transformer [36] is used as a branch of CircSite model, it mainly consists of positional encoding, multi-head attention, layer normalization, and MLP. It focuses on learning global information buried in circRNA sequences.

#### 2.5.1 Positional Encoding

In addition to word (a patch) embedding, the embedded patches in Fig. 1 need positional embedding to represent the position of the word in the sequence. The Transformer does not use the structure of an RNN, and cannot make use of the order information of the words in a sequence. Thus, position embedding (PE) is used in Transformer to preserve the relative or absolute position of words in a sequence. PE has the same dimension as word embedding and can be obtained by either training or predefining. In this study, PE is defined as below:

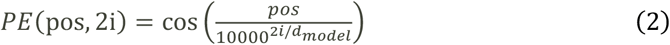

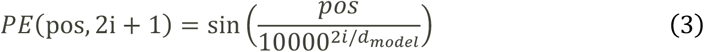

where pos denotes the position of the word (patch) in the sequence, *d* denotes the dimension of the *PE*, 2*i* denotes the even index, and 2*i* + 1 denotes the odd index (i.e., 2i ≤ d, 2i + 1 ≤ d).

#### 2.5.2 Multi-Head Attention

Self-attention takes the representation of a word in three matrices *W*_*Q*_, *W*_*K*_, *W*_*V*_ to obtain three attention values Q, K and V. Self-attention is calculated as below:

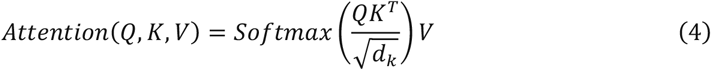

where *d*_*k*_ is the dimension of the matrix *K*.

Multi-Head attention is the attention calculated after *Q, K*, and *V* through h linear maps, where h is set to 12. Finally, all the outputs from *Q, K*, and *V* are concatenated to improve the learning ability and expressive power of the deep network.

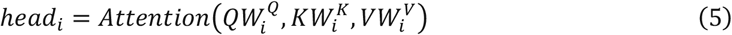

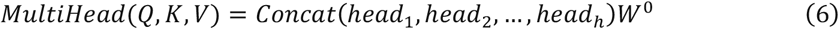

#### 2.5.3 Layer normalization

Layer normalization (LN) [37] is a batch size-independent algorithm, the number of samples in a batch does not affect the amount of data involved in the LN. The dimensions of the layer regularization operation are [*C, H, W*], where *C, H* and *W* is the channels, the height and width of feature maps. The mean and variance are normalized for each input of a layer of the network.

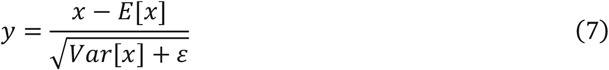

where *E* and *Var* are the mean and variance, respectively, and *ε* is a very small number to prevent the error that the denominator is zero.

#### 2.5.4 MLP layer

In the MLP layer of the encoder, we use two fully connected layers with activation function GELU.

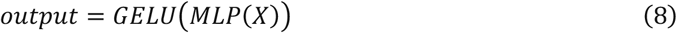

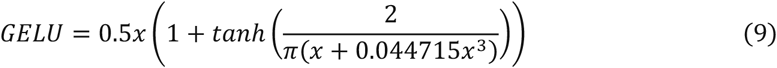

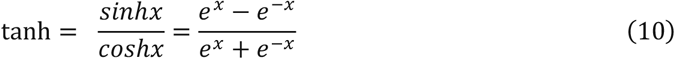

### 2.6. Concatenation layer

In this layer, we combine the representations learned from the BiGRU layer and the encoder of Transformer, and the combined representations are fed into the final MLP classifier.

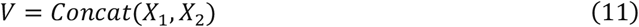

where *V* is the fused representations of the circRNA, *X*_1_ and *X*_2_ are the representations learned by BiGRU and Transformer, respectively.

### 2.7. MLP classification layer

The MLP layer consists of a fully connected layer with the activation function Sigmoid:

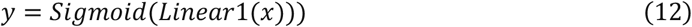

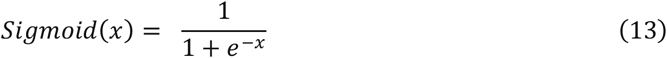

### 2.8. Loss function

In this study, predicting RBP binding nucleotides on RNAs is formulated as a binary classification problem, and the output score of the model is the probability of the centering nucleotide interacting with the RBP. The loss function is defined as the sum of the binary cross-entropy loss and the *l*_2_ regularization term acting as a weight decay to prevent the model from overfitting:

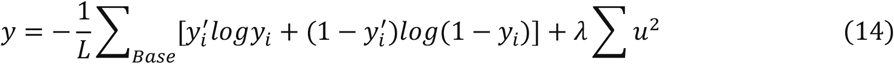

Where 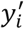, *y*_*i*_ denotes the true label and predicted score, respectively, *λ* is a hyperparameter of *l*_2_ regularization term penalty weight, and *u* is the learnable parameters of the model.

### 2.9. Integrated gradient for detecting key subsequences

To demonstrate the power of the trained deep model, we use Integrated Gradient (IG) [31] to calculate the contribution score of individual nucleotides on the sequences. IG solves the gradient saturation of the traditional gradient-based interpretability method. For example, the elephant’s trunk is very important for the algorithm’s decision to recognize an object as an elephant. However, when the length of the elephant’s trunk increases to a certain extent, the decision score will not increase, resulting in a gradient of 0 between the output and the input feature. In this study, we change the size of input features through linear interpolation, and then calculate the back-propagation gradient of each feature. The larger the gradient is, the more important the feature is. An important operation of IG is that it does not draw a conclusion only based on one interpolation, but based on the integral value of the gradients within the interpolation range:

1. The input of CircSite is a one-hot encoding matrix with only two types of values (0 and 1). We divide 1 equally into 100 bins, thus, 100 interpolation matrices are obtained. Theoretically, more interpolation is more accurate to calculate the gradient change, but it will undoubtedly increase the computational burden. We experimentally find that 100 interpolation bins are enough to draw an accurate gradient change curve.
2. We first fed the interpolation matrix into the trained deep model and calculate the change in gradient after the change in the size of each feature.
3. Then we obtain the importance of a feature based on the integral value of the gradient line of the 100 interpolation matrices.

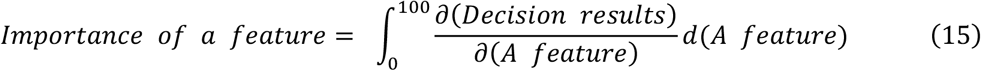

### 2.10. Score binarization

Generally, one RBP binding site consists of several consecutive nucleotides. Our goal is not only to predict the interaction score of each nucleotide on the circRNAs, but also to decide which nucleotides are located in the true binding sites. Thus, we binarize the raw predicted scores to determine the boundary of the binding site.

First, we process the raw predicted scores on the circRNAs with median filtering to leverage binding status of neighboring nucelotides, then use the threshold determined on the validation set to detect a candidate binding region instead of the default threshold 0.5 for all RBPs. We count the range of circRNA regions interacting with RBP in our benchmark datasets and find that the minimum range is 23. Thus, we finally choose 23 as the size of the candidate binding regions of RBPs. If the length of the candidate region is greater than 23, we consider this region to be the binding region and set all values of this region to 1, otherwise set it to 0. The labels of individual nucleotides are obtained using a novel scoring binarization strategy, whose pseudo-code is given in Algorithm 1.

#### Algorithm 1

Score binarization strategy

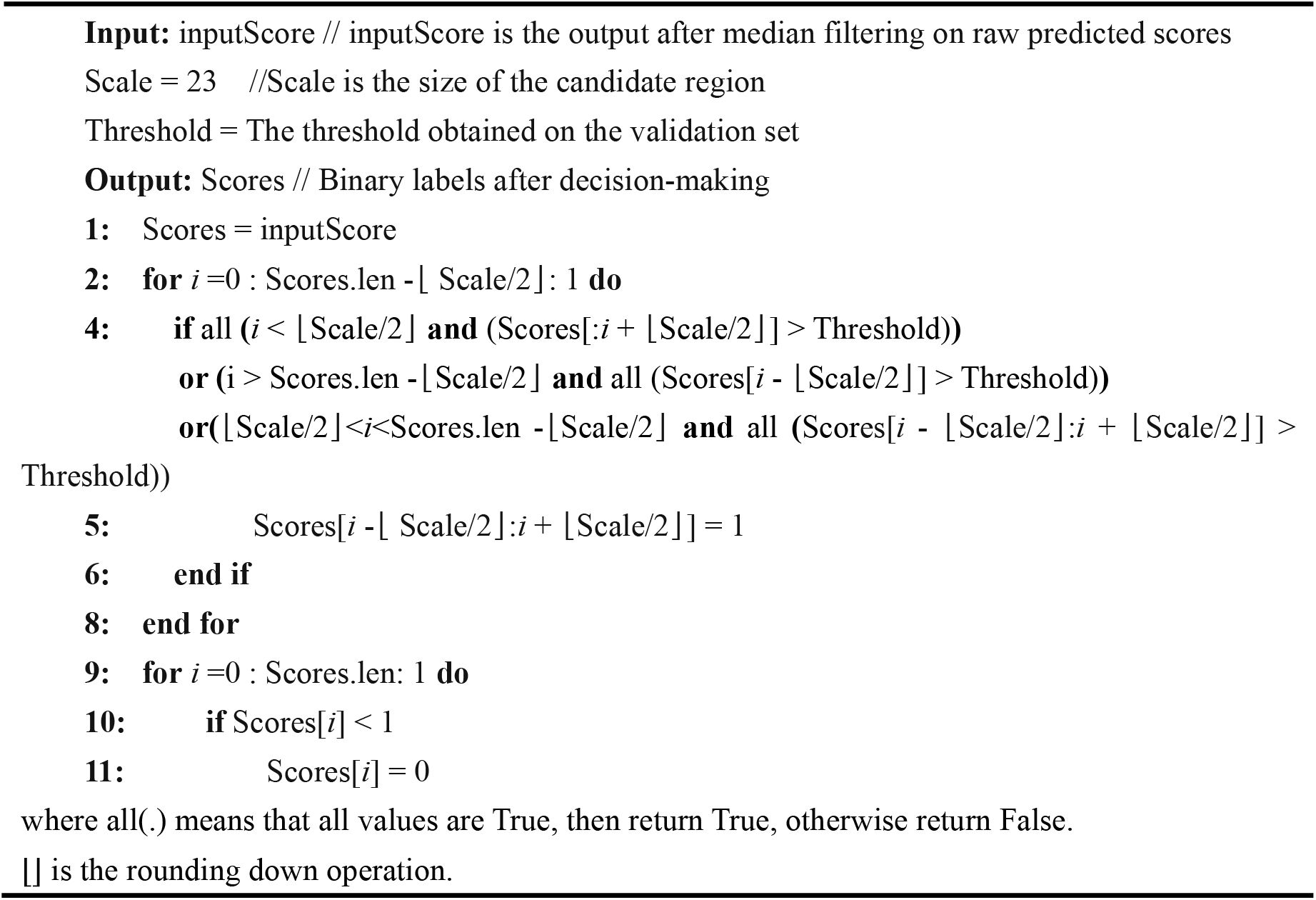

### 2.11. Evaluation metrics

In this study, we objectively evaluate the CircSite from three perspectives: nucleotide-level predictions, region-level predictions and whole-level predictions. The former two metrics are calculated on the nucleotide-level test set and the latter one is calculated on the independent circRNA test set.

#### 2.11.1 Nucleotide-level metrics

Each circRNA is split into fragments centering at the nucleotides using a sliding window, which results in an imbalanced test set. Here the scores of individual fragments are treated as the binding scores of individual nucleotides centering at the fragments along the circRNAs. In this study, area under precision-recall curve (auPRC) and area under receive operative curve (auROC) are used as the performance metric. Due to data imbalance between positive and negative samples in the test set, we focus on the auPRC when comparing different methods. After median filtering and score binarization on the raw scores, Accuracy, Matthews Correlation Coefficient (MCC), Precision, Recall and F1 score are calculated for nucleotide-level prediction where each nucleotide is a sample.

#### 2.11.2 Region-level metrics

We evaluate CircSite for region-level prediction. Binding regions are defined as a fragment consisting of several consecutive nucleotides. First, we binarized the prediction scores according to a threshold derived from our score binarization strategy. Several consecutive binding nucleotides (over 23) are treated as a binding region. A prediction is considered correct if it satisfies the condition that the overlap between the true binding region and the predicted binding region is at least half of the longer region. Finally, we calculate the following metrics for region-level prediction where each region is a sample:

Precision based on interaction regions (*PRE*_*B*_):

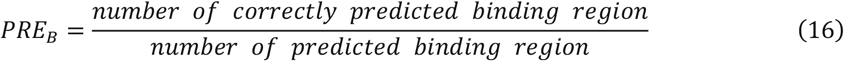

Recall based on interaction regions (*REC*_*B*_):

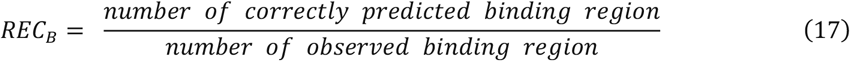

The *F*1_*B*_ score is based on the binding region, where *F*_1_ is a metric that takes into account both the accuracy and recall of the classification model:

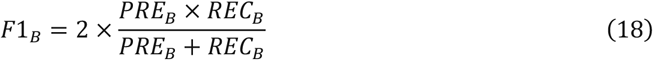

#### 2.11.3 Whole-level metrics

To assess the overall ability of a circRNA interacting with a given RBP, we evaluate whether CircSite is able to predict a circRNA binds to a given RBP. For circRNAs, the binding scores of individual nucleotides are predicted for a given RBP, followed by median filtering and score binarization. If a circRNA has at least one binding region that can interact with the given RBP, then the prediction for this circRNA is 1, otherwise 0. Then, Accuracy, Matthews Correlation Coefficient (MCC), Precision, Recall, and F1 score are calculated as whole-level metrics where each RNA is a sample.

### 2.12. Experimental settings

In CircSite, we first map each subsequence of a length 7 into an embedding using 384 1-D convolutional layers with convolutional kernels of a size 7, then the embedding is fed into the BiGRU module and the Transformer module, respectively. The BiGRU module consists of 2 layers GRUs with 384 hidden neurons in both directions. The Transformer module uses 12 layers as the block, it consists of a query vector, a key vector, and a value vector, i.e., the parameter matrix of *W*^*Q*^, *W*^*K*^, *W*^*V*^, all the vectors have a dimension of 384. Next, we obtain the representations by combining the two branches, and the MLP classification layer uses a fully connected layer of a dimension [384, 1] with a Sigmoid activation function. The optimizer of the model is Adam with a learning rate of 1e-4, and the loss function is a binary cross-entropy loss function. Early stopping is used to mitigate the risk of model overfitting. All the above parameters are determined by cross-validation approach. Details of the parameters are given in Supplementary Table S3.

### 2.13. Baseline methods

To evaluate the nucleotide-level prediction performance of CircSite, we compare it with four baseline models as follows:

1. **Random Classifier**: It randomly generate scores between 0-1 for test samples.
2. **CNN-BiGRU**: An 1-D convolution layer with 384 convolutional kernels of a kernel size 7 and step size 1 is used to map each RNA sequence of a length 7 into a patch embedding, which is fed into a bidirectional BiGRU of a hidden size 384, and the output of the last time step is fed into a fully connected layer with Sigmoid activation function. The hybrid CNN and LSTM are also used in iDeepS and DeepCLIP.
3. **Transformer**: The encoder part of the Transformer is used, both the number of multi-heads and depth are set to 12. The output of the 1-D convolution layer with the positional encoding is fed into the Transformer to make the binary classification.
4. **iCircRBP-DHN**: It uses a deep hierarchical network with attention mechanism from multiple sources of features to classify RBP binding fragments of circRNAs, iCircRBP-DHN has been shown to be superior to the CRIP method [14] on predicting whether a circRNA fragment contains a binding site or not. Thus, we do not compare CircSite with CRIP. Here, we modify iCircRBP-DHN to score the fragment centering at a nucleotide as the binding score of this nucleotide, then obtain the binding scores for all nucleotides on circRNAs. To evaluate the whole-level prediction performance of CircSite, we compare it with catRAPID omics v2.0 [38] using proteins and circRNA sequences as the input:
5. **catRAPID omics v2.0** [38]: It divides the circRNA into fragments and calculates the RBP-circRNA interaction ranking scores based on physico-chemical profiles [39] using a precompiled RBP or circRNA library. The ranking score is scaled to [0, 1] and consists of normalized interaction propensity, RBP propensity and the presence of known RNA-binding motif.

## 3. Results

### 3.1. CircSite achieves promising performance on nucleotide-level prediction

In this section, we compare CircSite with the Random classifier, CNN-BiGRU, and Transformer, on predicting RBP binding nucleotides on circRNAs in the nucleotide-level circRNAs test set, where all four methods are trained on the same nucleotide-level training set. Here we treat each nucleotide on the circRNAs as a sample when calculating the performance metrics.

We first optimize the size of the sliding window for CircSite to achieve the best auPRC on validation sets. We analyze the RBP binding sites on circRNAs and find that over 90% of the circRNAs interacting with RBPs have binding sites that have less than 100 nucleotides. Thus, we use grid search to evaluate the window size with the range of 25-85 with a step five on the validation sets of five randomly selected RBP datasets. As shown in Supplementary Fig. S1, CircSite yields the highest average auPRC on the five validation sets. Thus, we finally choose 65 as the sliding window size for CircSite for the following experiments.

Next, we compare CircSite with its individual components on the nucleotide-level test set. As shown in Fig. 2A and 2B, CircSite is superior to other three methods (Supplementary Table S4). For the 37 RBPs, CircSite consisting of hybrid BiGRU and Transformer achieves an average auPRC of 0.3292 and an average auROC of 0.7809. Since the test set is extremely imbalanced, we are more focused on auPRC when comparing different methods. Compared with the random classifier baseline, CircSite yields over tenfold improvement on auPRC (auPRC of 0.0382, Wilcoxon rank-sum test *P*-value of 9.811e-14). CircSite achieves a relative increase of about 60% on auPRC compared to Transformer (auPRC of 0.2008, Wilcoxon rank-sum test *P*-value of 3.799e-5) and a relative increase of about 10% over an auPRC of 0.2988 (Wilcoxon rank-sum test *P*-value of 0.152) of CNN-BiGRU. Of the 37 RBPs, CircSite achieves higher auPRC than Transformer and CNN-BiGRU on 37 and 34 RBPs, respectively. As shown in Supplementary Table S4, we can see that Transformer yields a higher auROC, but CNN-BiGRU obtains a higher auPRC. The results indicate that Transformer and CNN-BiGRU can complement each other. Thus, CircSite integrates Transformer and BiGRU to improve the prediction performance. For some RBPs, CircSite yields a large increase in auPRC, especially for those RBPs with a small number of binding targets, demonstrating the effectiveness of the nucleotide-level training set construction strategy. For the RBP WTAP, CircSite increases the auPRC 0.1565 of CNN-BiGRU to 0.2825, a relative increase of 81%.

**Fig. 2.**
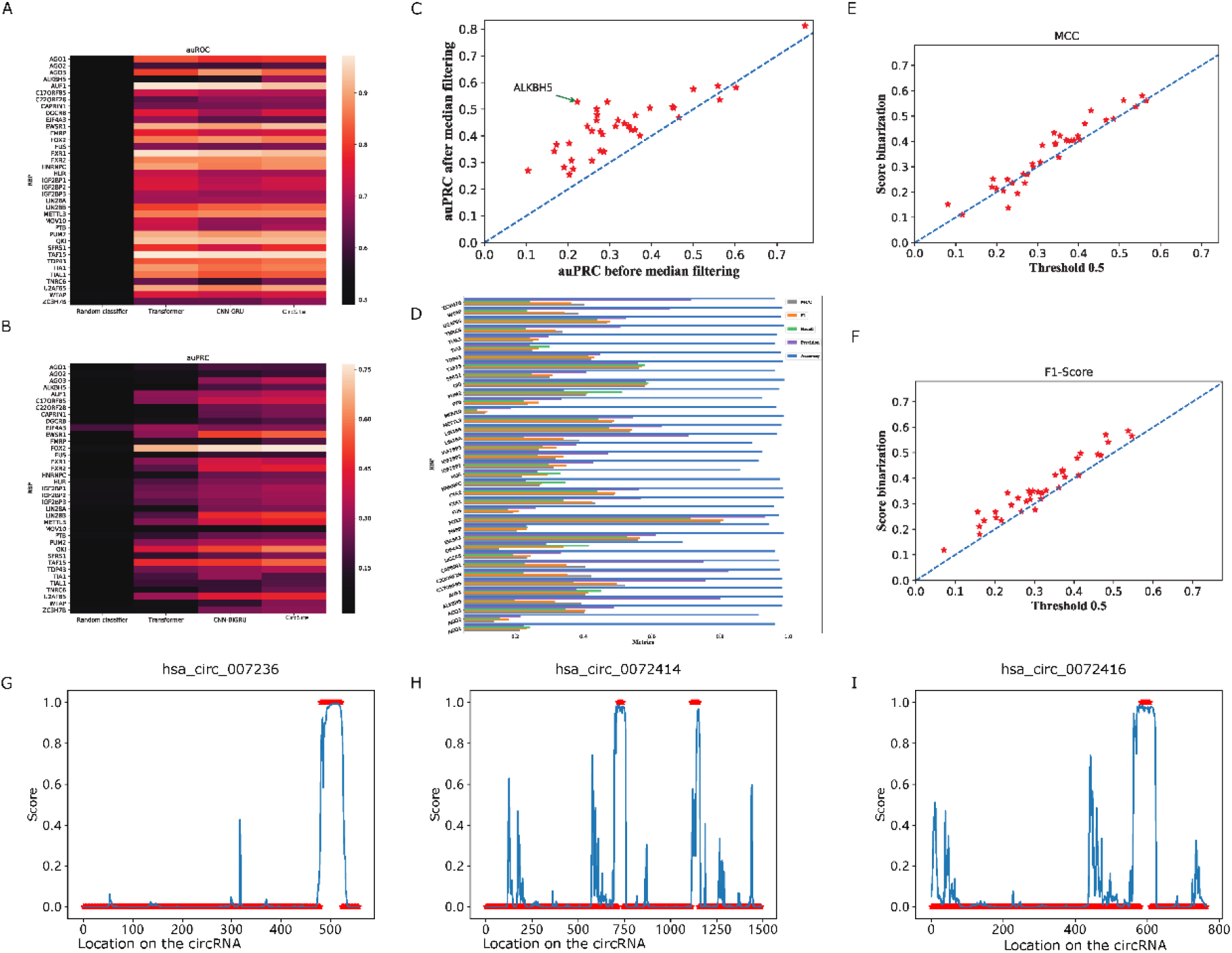
Performance comparison of CircSite and CNN-BiGRU, Transformer and Random Classifier on the nucleotide-level test set. A) auROC based on raw predicted scores of individual nucleotides by MLP classification layer in CircSite; B) auPRC based on raw predicted scores of individual nucleotides by MLP classification layer in CircSite and C) auPRC comparison of CircSite with and without median filtering. D) MCC, F1-score, Recall, Precision and Accuracy metrics based on the final binary values determined by CircSite. E) MCC comparison of CircSite with the score binarization and the default threshold 0.5. F) F1-score comparison of CircSite with the score binarization and the default threshold 0.5. G, H and I are the raw scores of individual nucleotides on hsa_circ_0072336, hsa_circ_0072414 and hsa_circ_0072416 interacting with AGO1, respectively. Red points are the ground truth and blue points are the predicted scores by the MLP classification layer in CircSite.

Furthermore, we apply median filtering to leverage the binding information of neighboring nucleotides, further smoothing the raw predicted scores to remove the false positive binding nucleotides. To demonstrate the added value of median filtering, we compare the auPRC of CircSite with and without median filter. As shown in Fig. 2C and Supplementary Table S5, we can see that median filter improves the average auPRC across 37 RBPs from 0.3292 to 0.4382 (a relative increase of 33.11%), indicating that median filtering removes many false positives given that there exist extremely data imbalance in the test set. The results demonstrate that it is necessary to integrate the median filtering module into CircSite. In addition, we report other nucleotide-level metrics (Accuracy, Precision, Recall, F1-score, and MCC) after score binarization for CircSite. They are evaluated on the nucleotides in the nucleotide-level test set, where the binding scores are binarized according to the threshold obtained on the validation set of different RBPs. As shown in Fig. 2D, CircSite yields an accuracy of 0.9553, precision of 0.4782, recall of 0.3318, F1-score of 0.3721 and MCC of 0.3644, respectively. We can see that different threshold values are used for different RBPs to determine a binding nucleotide; they range from 0.09 of FUS to 0.86 of FOX2 (Supplementary Table S6). We further demonstrate the added value of score binarization (Supplementary Table S7). CircSite with the default threshold 0.5 yields an accuracy of 0.9323, precision of 0.5222, recall of 0.2707, F1-score of 0.343 and MCC of 0.3289, respectively. We can see that the objective metrics MCC (Fig. 2E) and F1-score (Fig. 2F) are on average both lower than those of CircSite with our proposed score binarization. The results show that it is necessary to choose different threshold values for score binarization on the validation sets of RBPs instead of using the default threshold 0.5.

Additionally, we visualize the raw prediction scores of the MLP classification layer in CircSite along the circRNA, we test three randomly selected circRNAs using the trained model of AGO1. As shown in Fig. 2G-I, we can see that CircSite outputs high scores for those true binding sites. For the four circRNAs, CircSite detects all the binding sites, including the has_circ_0072414 with two true binding sites. However, some unbound regions have high predicted raw scores, which may result in false binding nucleotides. The results demonstrate that CircSite is able to detect the true binding nucleotides accurately, but it is still necessary to remove those unbinding regions with high predicted scores.

We further illustrate the effectiveness of median filtering and novel score binarization strategy to post-process the raw scores, further removing the false binding nucleotides. Supplementary Fig. S2A and D show the raw predicted scores of nucleotides on the two circRNAs with the RBP ZC3H7B without any post-processing. Based on the signal maps, a median filter with a window size 46 in CircSite is able to smooth the scores and further decrease the scores for the non-binding nucleotides (Supplementary Fig. S2B and E). Furthermore, the new score binarization strategy in CircSite transforms the raw scores to be 0 or 1 as decision-making results (Supplementary Fig. S2C and F). The results demonstrate that median filtering and new score binarization strategy are able to potentially reduce the number of false binding nucleotides.

### 3.2. CircSite is superior to the state-of-the-art method on nucleotide-level prediction

We compare CircSite with the state-of-the-art method iCircRBP-DHN designed for inferring RBP-binding circRNA fragments and we modify iCircRBP-DHN to predict binding nucleotides on circRNAs. Here, we train two types of iCircRBP-DHN models: 1) iCircRBP-DHN^1^ is trained on original circRNA fragments and evaluated on our nucleotide-level test set. 2) iCircRBP-DHN^2^ is trained and evaluated on our nucleotide-level training and test set.

As shown in Fig. 3A, iCircRBP-DHN^1^ trained on original circRNAs fragments yields relatively poor results (an average auROC of 0.637 and auRPC of 0.0808 across 37 datasets) on our nucleotide-level test set, which are much worse than CircSite with an average auROC of 0.7890 and auPRC of 0.3292, and the Wilcoxon rank-sum test *P*-value for auPRC comparison is 7.056e-12. CircSite outperforms iCircRBP-DHN^1^ on all 37 datasets and the detailed results are given in Supplementary Table S8. One potential reason is that iCircRBP-DHN^1^ is designed for circRNA fragments, it predicts whether fragments bind to an RBP or not. It is not capable of capturing contextual information for predicting RBP-binding nucleotides, although iCircRBP-DHN^1^ yields high accuracy on the original fragment test set. In addition, the balanced fragment-based training set and extremely imbalanced nucleotide-level test set are differently distributed, and the trained model on original fragments cannot capture patterns of negative samples and better discriminate the patterns of positive and negative samples. The results demonstrate that the deep model trained on original circRNA fragments cannot generalize well to nucleotides on circRNAs and constructing a new training set for RBP-binding nucleotides is necessary.

**Fig 3.**
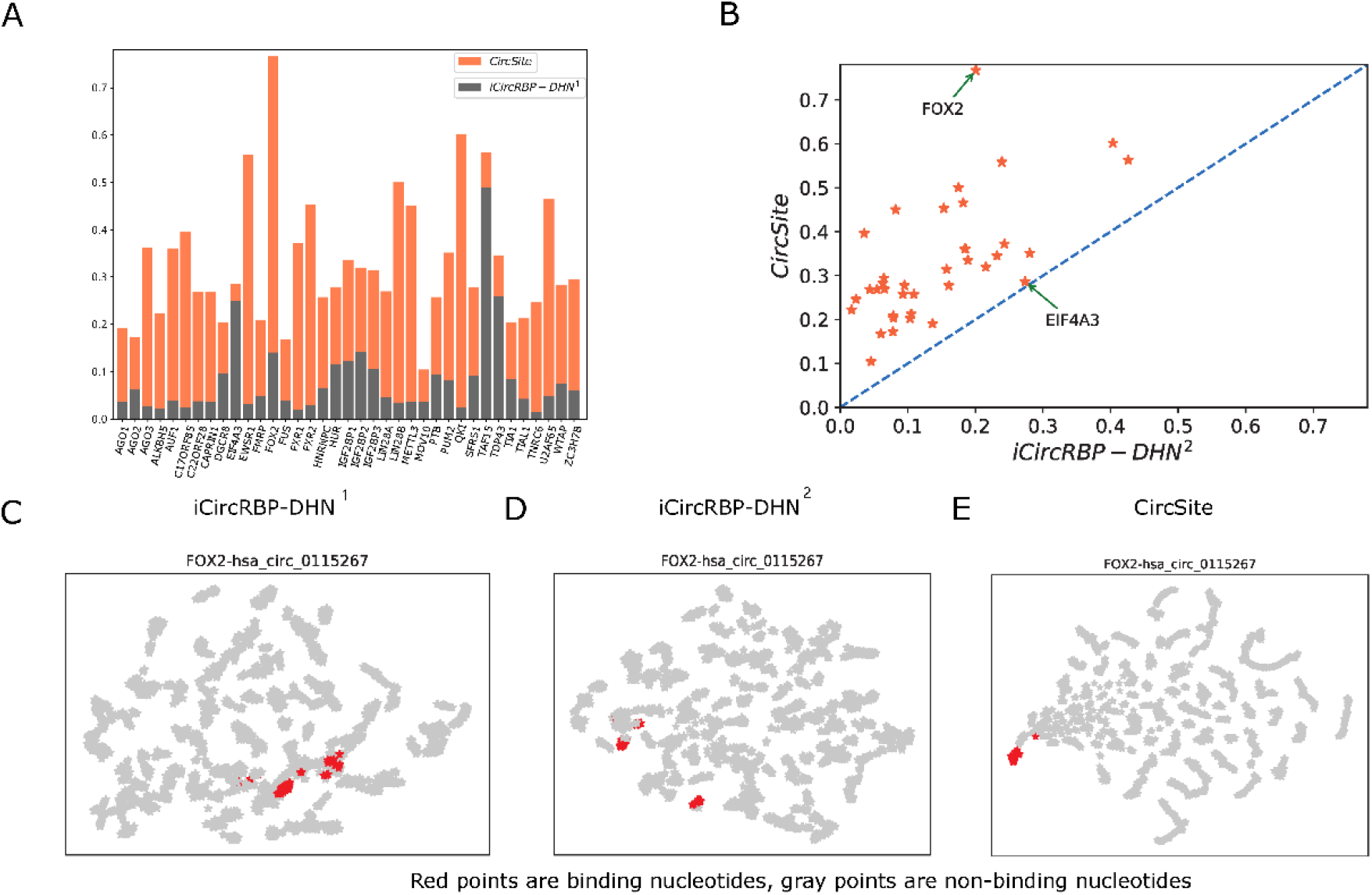
AuPRC comparison between CircSite and the modified iCircRBP-DHN for predicting binding nucleotides. A) CircSite is trained and tested on our nucleotide-level dataset and iCircRBP-DHN^1^ is trained on the training set of iCircRBP-DHN and evaluated on our nucleotide-level test set. B) CircSite and iCircRBP-DHN^2^ are trained and evaluated on our nucleotide-level training and test set. C, D, E are the 2-D visualization of nucleotides on has_circ_0115267 binding to FOX2 learned by iCircRBP-DHN^1^, iCircRBP-DHN^2^ and CircSite.

To demonstrate the effectiveness of our training set construction strategy, we train the iCircRBP-DHN^2^ method on our nucleotide-level training and test set. We can see that the performance is improved with a large margin (Supplementary Table S9) compared to iCircRBP-DHN^1^. It increases the average auROC from 0.6370 to 0.7361 and average auPRC from 0.0808 to 0.1442, and the Wilcoxon rank-sum test *P*-value for auPRC comparison is 3.894e-9. The results indicate that our constructed nucleotide-level training set can capture more contextual information for predicting RBP-binding nucleotides. Furthermore, we compare CircSite with iCircRBP-DHN^2^ on the same nucleotide-level training and test set. As shown in Fig. 3B, the auPRCs of CircSite are higher than iCircRBP-DHN^2^ for all 37 RBPs. For FOX2, CircSite yields an auPRC of 0.7672, which is over three times than 0.2007 of iCircRBP-DHN^2^. The results demonstrate that the designed hybrid Transformer and BiGRU model in CircSite is superior to the deep multi-scale residual network in iCircRBP-DHN. It is able to capture discriminative contextual information for RBP-binding nucleotides prediction.

Moreover, we illustrate the learned embeddings of the nucleotides on the has_circ_0115267 binding to FOX2 by iCircRBP-DHN^1^, iCircRBP-DHN^2^ and CircSite in 2-D space using T-SNE [40], where the number of binding nucleotides are much fewer than the number of non-binding nucleotides. As shown in Fig. 3C-E, the binding nucleotides are better separated from the non-binding nucleotides in the 2-D embedding space of CircSite than iCircRBP-DHN^1^ and iCircRBP-DHN^2^. We can see that the binding nucleotides are mixed with non-binding nucleotides in the 2-D embedding space of iCircRBP-DHN. The results demonstrate that CircSite is able to learn more discriminate emebddings for predicting RBP-binding nucleotides than iCircRBP-DHN.

Furthermore, we use TAF15 as an example to illustrate the results obtained in iCircRBP-DHN^1^, iCircRBP-DHN^2^ and CircSite on two circRNAs. As shown in Supplementary Fig. S3, iCircRBP-DHN^1^ predicts most of the nucleotides as the binding nucleotides, which result in too many false positives. As mentioned above, the potential reason is that the distribution of positive and negative samples in the training set is different from those in the test set, resulting in the covariate shift. Differently, iCircRBP-DHN^2^ and CircSite is able to remove most of the false positives, considering that they are trained the same nucleotide-level training and test set with a similar ratio of positive and negative samples. On the other hand, we can see that CircSite predicts much fewer false positives than iCircRBP-DHN^2^, demonstrating the effectiveness of CircSite for predicting RBP-binding nucleotides.

### 3.3. CircSite achieves promising performance on region-level prediction

Since the binding sites corresponding to binding regions may consist of multiple binding nucleotides, we treat each region as a sample and evaluate CircSite for predicting RBP-binding regions. The predicted binding regions are obtained after median filtering and score binarization in CircSite to predict RBP-binding regions, and we calculate *PRE*_*B*_, *REC*_*R*_, and *F*1_*B*_ score as performance metrics. Here we treat each binding region as a sample when calculating the performance metrics.

In this study, we calculate two types of the region-based metrics according to two definitions of correct predictions and the details of the results are shown in Fig. 4 and Supplementary Table S10. When we define the prediction as correct if there exists overlap between true and predicted binding region, for the 37 RBPs, CircSite yields an average 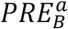 of 0.5664, 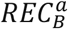of 0.3717 and 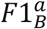 score of 0.4243, respectively. Furthermore, we define the correct prediction more strictly when the overlap between true and predicted region overlap at least 50% of the larger region, CircSite yields an average 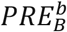 of 0.4950, 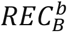 of 0.3075 and 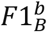 score of 0.3592, respectively. We can see that the strict definition of correct predicted binding region decrease to a certain extent the prediction performance (Fig. 4). However, CircSite still yields a precision close to 0.5 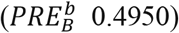. The results demonstrate that CircSite is able to predict RBP-binding regions with promising performance.

**Fig. 4.**
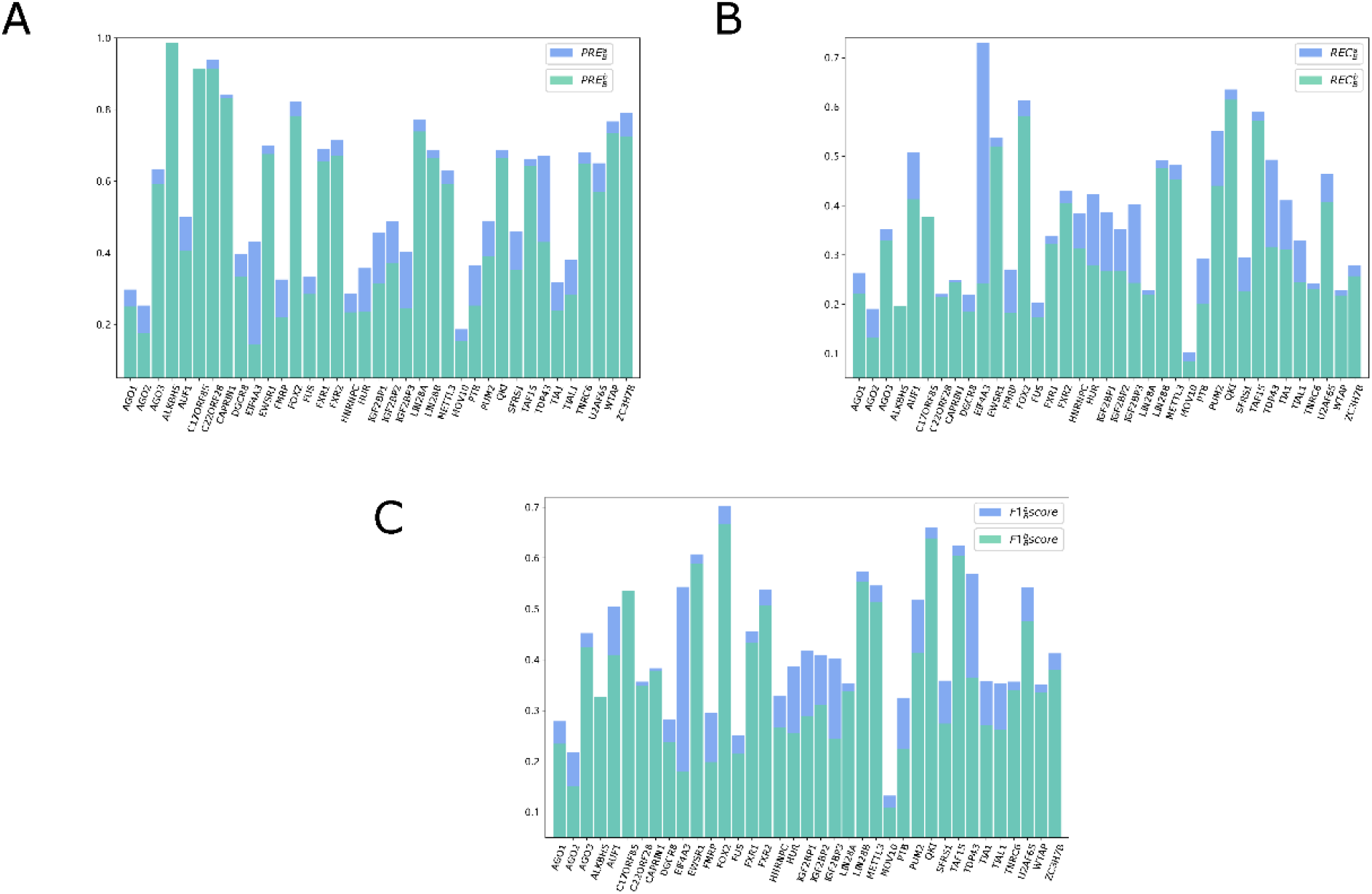
Region-based metrics based on two definitions for correct predicted regions. a: The prediction is considered correct if the true binding region overlaps with the predicted region; b: The prediction is considered correct if the overlapped region between the true binding region and the predicted region takes over 50% of the largest region. A) Precision comparison of two definitions; B) Recall comparison of two definitions; C) F1 score comparison of two definitions.

### 3.4. CircSite enables inferring RBP-binding circRNAs

To further explore the power of CircSite, we construct 37 RBP independent test sets to predict whether a circRNA can interact with an RBP, here we treat each circRNA as a sample when calculating the performance metrics. We evaluate CircSite and catRAPID omics v2.0 on the same test sets, and we upload the RBP sequences and circRNA sequences to catRAPID omics v2.0 webserver to obtain the ranking scores, the default threshold 0.5 for catRAPID omics v2.0 is used to make a binary decision.

As shown in Supplementary Table 11 and Fig. 5, CircSite yields an average accuracy of 0.716, MCC of 0.461, precision of 0.813, recall of 0.595, and F1-score of 0.668 across 37 RBPs. The accuracy ranges from 0.615 for LIN28A to 0.841 for HNRNPC on the balanced independent test set. However, catRAPID omics v2.0 achieves an average accuracy of 0.543, MCC of 0.115, precision of 0.536, recall of 0.521, and F1-score of 0.452 across 37 RBPs. All the average metrics of catRAPID omics v2.0 are lower than those of CircSite (Fig. 5), especially for the objective metric MCC. Of the 37 RBPs, CircSite achieves an accuracy over 0.7 for 20 RBPs, outperforming catRAPID omics v2.0 for 36 RBPs. The potential reasons are due to the followings: 1) catRAPID omics v2.0 is a general model trained on pooled RNA-RBP interactions from different RBPs. The binding patterns of individual RBPs are different, introducing contradictory binding patterns for general models. However, CircSite trains a model per RBP, it is able to train a more accurate and specific model when a large volume of binding data is available for each RBP. 2) catRAPID omics v2.0 is designed for multiple types of coding and non-coding RNAs, whose binding patterns may be different from circRNAs. 3) The threshold 0.5 is not optimized for catRAPID omics v2.0 to make a binary classification. The results show that CircSite is superior to catRAPID omics v2.0 and effective in predicting whether the circRNAs interact with RBPs or not, enabling inferring RBP-binding circRNAs.

**Fig. 5.**
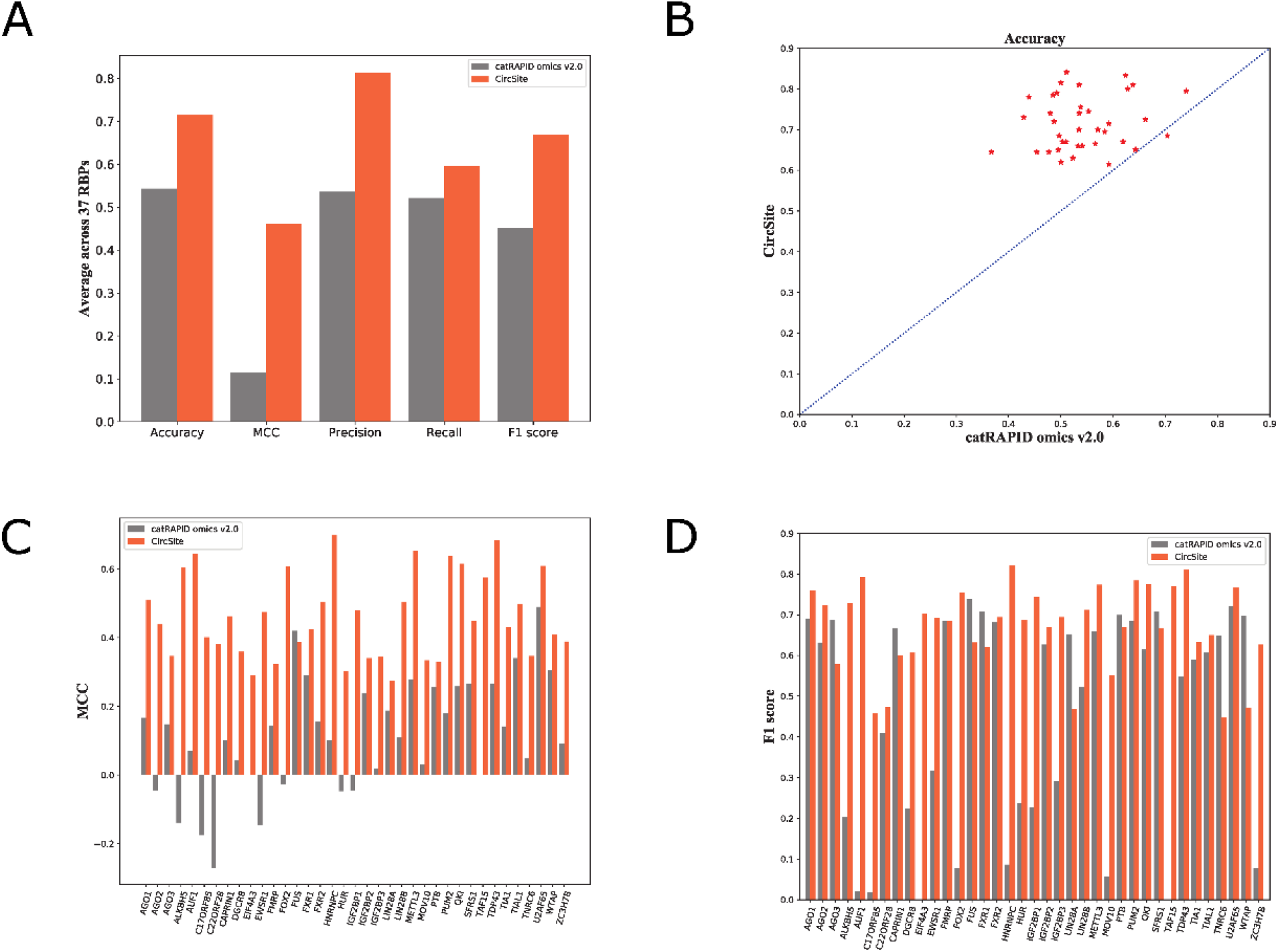
Performance comparison between CircSite and catRAPID omics v2.0 for predicting RBP-binding circRNAs on independent circRNA test set. A) The average metrics across 37 RBPs; B) Accuracy; C) MCC values; D) F1-score.

### 3.5. CircSite is able to explore the key subsequences aligning well with binding motifs on circRNAs

To further demonstrate the trained deep model is able to capture true binding patterns of RBPs, we use integrate gradient to calculate the importance of individual nucleotides on the circRNAs, further revealing multiple binding motifs with some dissimilarities for the same RBP. We visually inspect the motifs detected by CircSite against the known motifs in the CISBP-RNA database [41]. Compared to previous tools CRIP and iCircRBP-DHN that cannot detect binding motifs from the trained models, the advantage of CircSite is able to infer multiple known binding motifs on the circRNAs from the trained model.

Fig. 6 visualizes the importance of individual nucleotides on the circRNAs for four RBPs. We can see that there are some important regions across the RNAs, and these important regions are visually similar to the known motifs of the RBP in CISBP-RNA database. For example, TIA1 has a preference for U-rich binding sites [42]. CircSite detects four U-rich regions on the hsa_circ_0084450. QKI prefers to bind to AC-rich regions on RNAs [41]. CircSite also detects four regions with rich AC on hsa_circ_0099821 and they are visually similar to the known motifs in CISBP-RNA database. For other two RBPs, we can also observe similar results that CircSite is able to capture the known motifs on the circRNAs. In addition, we can see that each circRNA has multiple detected binding motifs with some dissimilarity, CircSite reveals differential binding motifs on the circRNAs for the same RBP. The results demonstrate that CircSite is able to detect important regions on circRNAs and these important regions are consistent with the known binding motifs of the RBPs.

**Fig. 6.**
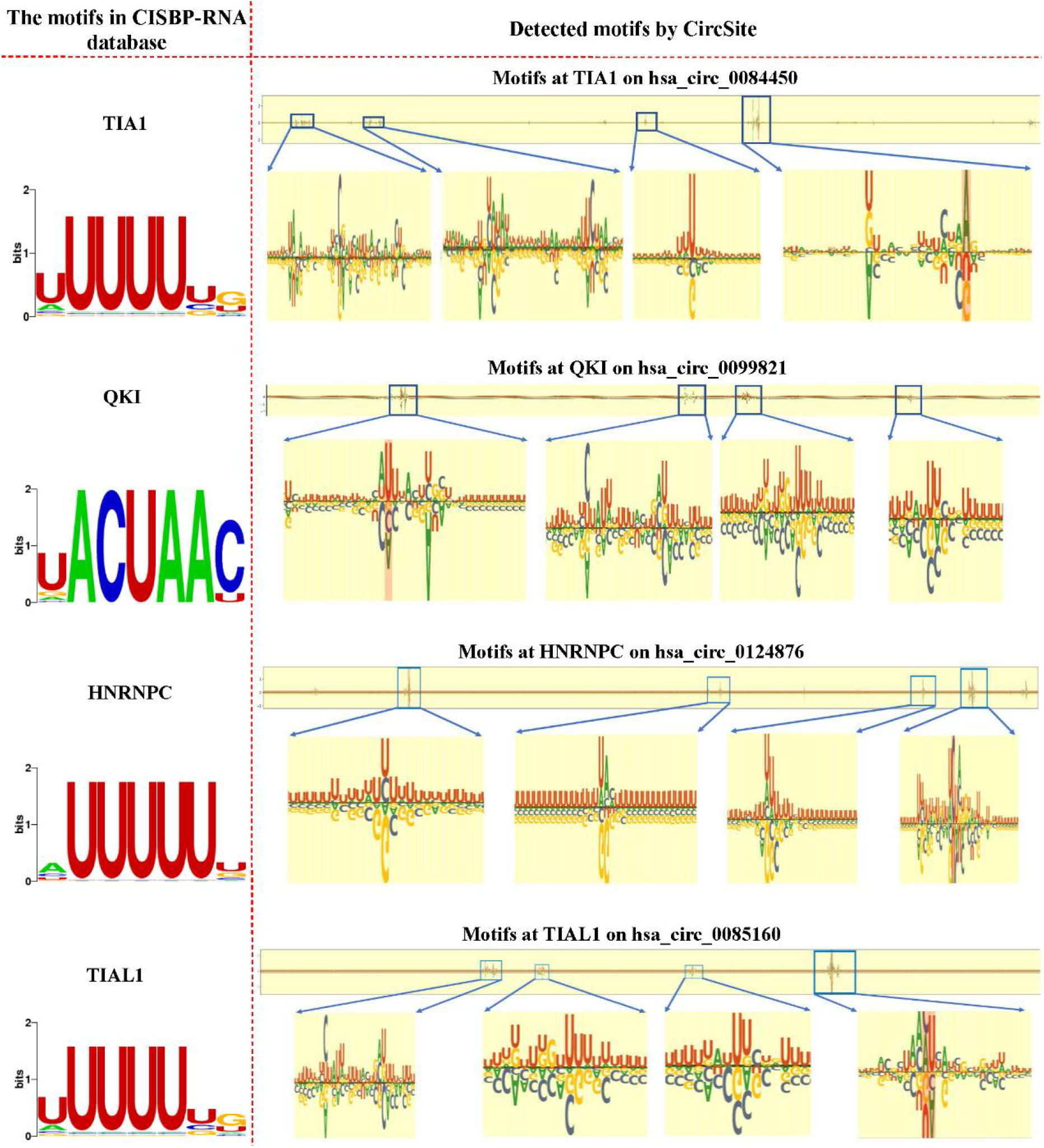
The importance of individual nucleotides on whole circRNAs by CircSite. The left side are the motifs obtained from the CISBP-RNA database, and the right side are the motifs detected by CircSite on whole circRNAs, larger positive values denote that nucleotides contribute more to predicting binding nucleotides.

## 4. Discussion

CircSite is able to predict RBP binding nucleotides on circRNAs, further inferring RBP binding labels for regions and RNAs. and CircSite is different from the existing methods for predicting RBP-RNA interactions. From short-read sequencing data, only the back-splice junction and the surrounding (about 100-300) nucleotides can be identified as part of a circRNA. Recently, a protocol is developed for circRNA sequencing using Oxford Nanopore Technology [43]. Previous methods construct the positive samples consisting of the binding fragments by extending upstream and downstream regions of the center of the binding site in a gene, and the same number of negative samples are extracted from the remaining fragments of the same gene. However, these methods can only predict whether the fragment contains a binding site or not. On the other hand, many non-binding fragments are discarded for model training, the methods cannot learn the patterns from more negative samples. Thus, many existing methods generally achieve high accuracy on the benchmark dataset. However, the false positive rate is very high for predicting binding nucleotides on circRNAs [29].

To resolve the above issue, CircSite uses a sliding window approach on the circRNAs to generate a large number of positive and negative fragments, whose labels are determined by the centering nucleotide of the fragment. All nucleotides on the circRNAs are involved in model training and evaluation. This strategy reduces the false positive rates to some extent, and makes CircSite be able to predict binding nucleotides on circRNAs. In addition, the circRNA sequences reported by CircInteractome are not experimentally validated sequences, but assumed sequences based on canonical splicing and host gene annotation. Thus, the CircSite model is built on presumed circRNA sequences, it should be rebuilt once there are more experimentally validated circRNA sequences available. Currently, our method is focused on circRNAs since there are binding circRNAs for a limited number of RBPs in Circinteractome database at this time. However, we can easily extend to other RBPs when more binding circRNA data are available in the future. In addition, we expect to extend our method to linear RNAs for more RBPs, i.e. ENCODE dataset.

Through the computational model CircSite, we can further explore its potential to discover that binding regions not detected as binding sites using high-throughput sequencing are predicted as potential binding sites. Take two RBPs, AUF1 and DGCR8, for example. As shown in Supplementary Fig. S4, the predicted binding regions are not experimentally verified, but the CircSite predicts that there is a high possibility that they are potential binding sites of RBPs, which will provide a useful guidance to follow-up biological experiments to verify these regions.

In contrast to existing RBP binding sites prediction methods, CircSite explores the effectiveness of a hybrid network model, median filtering, and new score binarization strategy to predict RBP binding nucleotides on circRNAs. The contributions of this study are:

1. A hybrid deep model is developed to improve the prediction performance by putting seven nucleotides as a patch and mapping them into a vector by convolutional layers, which are fed into BiGRU and Transformer, respectively. The local representations learned by CNN-BiGRU and the global representations learned by Transformer are concatenated as input for predicting RBP binding nucleotides.
2. Median filtering is applied on the predicted binding scores of individual nucleotides on circRNAs to obtain a smooth map of the predicted signals, it leverages the binding status of neighboring nucleotides to refine the predicted binding scores of individual nucleotides. Then a new score binarization strategy is used to decide whether the nucleotide interacts with the RBP using optimized thresholds for individual RBPs instead of using the default threshold 0.5 for decision making.
3. Compared to iCircRBP-DHN and CRIP, the binding sequence motifs of RBPs are mined from the learned CircSite model using integral gradient and these detected motifs are aligned well with known binding motifs. In addition, CircSite reveals multiple binding motifs with some dissimilarities on the circRNAs for the same RBP.
4. CircSite can not only predict the precise RBP binding nucleotides on circRNAs, but also predict whether the circRNAs can interact with a given RBP.

## 5. Conclusion

In this study, we present an interpretable hybrid network-based approach CircSite to predict binding landscape of RBPs on circRNAs at a single-nucleotide resolution, further inferring RBP binding circRNA regions and circRNAs. By using a sliding window to generate positive and negative samples on circRNAs, CircSite integrates a hybrid BiGRU and Transformer to improve the prediction performance. Furthermore, CircSite uses a median filter and novel score binarization strategy to incorporate prior biological knowledge to reduce false positive rate on predicting binding nucleotides. The results demonstrate that CircSite achieves higher accuracy and resolution for predicting RBP binding nucleotides. Moreover, we apply integrate gradient to detect key contributed subsequences, which are aligned well with known binding motifs of RBPs. It is expected that the CircSite tool will play important roles in the circRNAs and their binding proteins through the mode of binding site predictions at a base resolution.

## Code and data availability

The online webserver and datasets used in this study are available at http://www.csbio.sjtu.edu.cn/bioinf/CircSite/.

## REFERENCES

1. Huang, A., et al., Circular RNA-protein interactions: functions, mechanisms, and identification. Theranostics, 2020. 10(8): p. 3503–3517.

2. Hansen, T.B., et al., Natural RNA circles function as efficient microRNA sponges. Nature, 2013. 495(7441): p. 384–8.

3. Nishimasu, H., et al., Crystal structure of Cas9 in complex with guide RNA and target DNA. Cell, 2014. 156(5): p. 935–49.

4. Segel, M., et al., Mammalian retrovirus-like protein PEG10 packages its own mRNA and can be pseudotyped for mRNA delivery. Science, 2021. 373(6557): p. 882-+.

5. Okholm, T.L.H., et al., Transcriptome-wide profiles of circular RNA and RNA-binding protein interactions reveal effects on circular RNA biogenesis and cancer pathway expression. Genome Med, 2020. 12(1): p. 112.

6. Verduci, L., et al., CircRNAs: role in human diseases and potential use as biomarkers. Cell Death & Disease, 2021. 12(5).

7. Van Nostrand, E.L., et al., Robust transcriptome-wide discovery of RNA-binding protein binding sites with enhanced CLIP (eCLIP). Nature Methods, 2016. 13(6): p. 508-+.

8. Gerstberger, S., M. Hafner, and T. Tuschl, A census of human RNA-binding proteins. Nat Rev Genet, 2014. 15(12): p. 829–45.

9. Dunham, I., et al., An integrated encyclopedia of DNA elements in the human genome. Nature, 2012. 489(7414): p. 57–74.

10. Alipanahi, B., et al., Predicting the sequence specificities of DNA- and RNA-binding proteins by deep learning. Nature Biotechnology, 2015. 33(8): p. 831-+.

11. Pan, X. and H.B. Shen, RNA-protein binding motifs mining with a new hybrid deep learning based cross-domain knowledge integration approach. BMC Bioinformatics, 2017. 18(1): p. 136.

12. Gronning, A.G.B., et al., DeepCLIP: predicting the effect of mutations on protein-RNA binding with deep learning. Nucleic Acids Res, 2020. 48(13): p. 7099–7118.

13. Pan, X.Y. and H.B. Shen, Predicting RNA-protein binding sites and motifs through combining local and global deep convolutional neural networks. Bioinformatics, 2018. 34(20): p. 3427–3436.

14. Zhang, K., et al., CRIP: predicting circRNA-RBP-binding sites using a codon-based encoding and hybrid deep neural networks. RNA, 2019. 25(12): p. 1604–1615.

15. Zhang, S., et al., A deep learning framework for modeling structural features of RNA-binding protein targets. Nucleic Acids Res, 2016. 44(4): p. e32.

16. Yang, Y., et al., iCircRBP-DHN: identification of circRNA-RBP interaction sites using deep hierarchical network. Brief Bioinform, 2020.

17. Pan, X., et al., RBPsuite: RNA-protein binding sites prediction suite based on deep learning. BMC Genomics, 2020. 21(1): p. 884.

18. Ghanbari, M. and U. Ohler, Deep neural networks for interpreting RNA-binding protein target preferences. Genome Res, 2020. 30(2): p. 214–226.

19. Uhl, M., et al., RNAProt: an efficient and feature-rich RNA binding protein binding site predictor. Gigascience, 2021. 10(8).

20. Trabelsi, A., M. Chaabane, and A. Ben-Hur, Comprehensive evaluation of deep learning architectures for prediction of DNA/RNA sequence binding specificities. Bioinformatics, 2019. 35(14): p. i269–i277.

21. Koo, P.K., et al., Global importance analysis: An interpretability method to quantify importance of genomic features in deep neural networks. PLoS Comput Biol, 2021. 17(5): p. e1008925.

22. Kazan, H., et al., RNAcontext: A New Method for Learning the Sequence and Structure Binding Preferences of RNA-Binding Proteins. Plos Computational Biology, 2010. 6(7).

23. Yuan, L.L. and Y. Yang, DeCban: Prediction of circRNA-RBP Interaction Sites by Using Double Embeddings and Cross-Branch Attention Networks. Frontiers in Genetics, 2021. 11.

24. Yu, H., et al., beRBP: binding estimation for human RNA-binding proteins. Nucleic Acids Res, 2019. 47(5): p. e26.

25. Maticzka, D., et al., GraphProt: modeling binding preferences of RNA-binding proteins. Genome Biology, 2014. 15(1).

26. Strazar, M., et al., Orthogonal matrix factorization enables integrative analysis of multiple RNA binding proteins. Bioinformatics, 2016. 32(10): p. 1527–35.

27. Pan, X., et al., Prediction of RNA-protein sequence and structure binding preferences using deep convolutional and recurrent neural networks. BMC Genomics, 2018. 19(1): p. 511.

28. Jia, C., et al., PASSION: an ensemble neural network approach for identifying the binding sites of RBPs on circRNAs. Bioinformatics, 2020. 36(15): p. 4276–4282.

29. Wu, H., et al., Recognizing binding sites of poorly characterized RNA-binding proteins on circular RNAs using attention Siamese network. Briefings in Bioinformatics, 2021. bbab279.

30. Maticzka, D., et al., GraphProt: modeling binding preferences of RNA-binding proteins. Genome Biol, 2014. 15(1): p. R17.

31. Sundararajan, M., A. Taly, and Q. Yan, Axiomatic Attribution for Deep Networks. Proceedings of International Conference on Machine Learning, 2017. 70: p. 3319–3328.

32. Dudekula, D.B., et al., CircInteractome: a web tool for exploring circular RNAs and their interacting proteins and microRNAs. RNA biology, 2016. 13(1): p. 34–42.

33. Fu, L., et al., CD-HIT: accelerated for clustering the next-generation sequencing data. Bioinformatics, 2012. 28(23): p. 3150–3152.

34. Lecun, Y., et al., Gradient-based learning applied to document recognition. Proceedings of the Ieee, 1998. 86(11): p. 2278–2324.

35. Cho, K., et al., Learning Phrase Representations using RNN Encoder-Decoder for Statistical Machine Translation. Proceedings of the 2014 Conference on Empirical Methods in Natural Language Processing, 2014: p. 1724–1734.

36. Vaswani, A., et al., Attention Is All You Need. 31st Conference on Neural Information Processing Systems, 2017. 2017: p. 6000–6010.

37. Ba, J.L., J.R. Kiros, and G.E. Hinton, Layer normalization. arXiv, 2016. arXiv:1607.06450.

38. Armaos, A., et al., catRAPID omics v2.0: going deeper and wider in the prediction of protein-RNA interactions. Nucleic Acids Res, 2021. 49(W1): p. W72–W79.

39. Agostini, F., et al., catRAPID omics: a web server for large-scale prediction of protein-RNA interactions. Bioinformatics, 2013. 29(22): p. 2928–30.

40. van der Maaten, L. and G.E. Hinton, Visualizing High-Dimensional Data Using t-SNE. Journal of Machine Learning Research, 2008. 9: p. 2579–2605.

41. Ray, D., et al., A compendium of RNA-binding motifs for decoding gene regulation. Nature, 2013. 499(7457): p. 172–7.

42. Forch, P., et al., The apoptosis-promoting factor TIA-1 is a regulator of alternative pre-mRNA splicing. Molecular Cell, 2000. 6(5): p. 1089–1098.

43. Rahimi, K., et al., Nanopore sequencing of brain-derived full-length circRNAs reveals circRNA-specific exon usage, intron retention and microexons. Nat Commun, 2021. 12(1): p. 4825.

